# Evolutionary gain and loss of a plant pattern-recognition receptor for HAMP recognition

**DOI:** 10.1101/2022.03.30.484633

**Authors:** Simon Snoeck, Bradley W. Abramson, Anthony G. K. Garcia, Ashley N. Egan, Todd P. Michael, Adam D. Steinbrenner

## Abstract

Pattern recognition receptors (PRR) recognize distinct pathogen and herbivore-associated molecular patterns (PAMPs and HAMPs) and mediate activation of immune responses, but the evolution of new PRR sensing functions is not well understood. We employed comparative genomics and functional analysis to define evolutionary events leading to the sensing of the peptide HAMP inceptin (In11) by the PRR Inceptin Receptor (INR). Existing and *de novo* genome assemblies revealed that the presence of a functional *INR* gene corresponded with In11 response across 55 million years (my) of legume evolution, and that In11 recognition is unique to the clade of Phaseoloid legumes. The *INR* loci of certain Phaseoloid and non-Phaseoloid species also contain diverse INR-like homologues, suggesting that the evolution of INR receptor function ∼28 mya occurred after an ancestral gene insertion ∼32 mya. Functional analysis of chimeric and ancestrally reconstructed receptors revealed that specific AA differences in the C1 leucine-rich repeat (LRR) domain and C2 intervening motif likely mediated gain of In11 recognition. In summary, we present a conceptual model for the evolution of a defined PRR function based on patterns of *INR* variation in legumes.

## Introduction

Plants use a combination of constitutive and inducible defensive traits to resist challenges by pathogens and herbivores. Activation of inducible defenses requires perception of the threat^1^, mediated by an innate immune system that uses germline-encoded pattern recognition receptors (PRRs) at the cell surface to recognize pathogen, herbivore, and damage-associated molecular patterns (PAMPs, HAMPs, and DAMPs)^2,3^. Well-characterized PRRs in the large receptor kinase (RK) family comprise an extracellular domain, a single-pass transmembrane motif, and an intracellular kinase domain^4^. Receptor-like proteins (RLPs) share structural similarities with RKs but lack a kinase domain^5,6^. The most common extracellular domain of RK/RLPs is a series of leucine-rich repeats (LRRs), which are known to mediate elicitor binding and co-receptor association^7–11^. LRR-RK/RLPs form a large gene family; the *Arabidopsis thaliana* genome contains about 223 LRR-RKs and 57 LRR-RLPs^5,6,8,12^, and the number of annotated RK/RLPs per genome varies across and between plant families^11,13^. Moreover, comparative genomic analyses of LRR-RK/RLPs involved in biotic interactions have revealed strong diversifying selection, lineage-specific expanded gene clusters and immune receptor repertoire variation both within and between species^6,14–17^.

Evidence is accumulating for diverse roles of RLPs as immune sensors^14,17^. Various LRR-RLPs from Arabidopsis, tomato, and cowpea have been shown to detect molecular patterns from fungi, bacteria, parasitic weeds and herbivores^3,14^. Cf-9, Cf-4 and Cf-2 interact with respective molecular patterns of *Cladosporium fulvum*, Avr9, Avr4 and Avr2 (via host protein Rcr3), and are at least restricted to the *Solanum* genus^18–24^. Arabidopsis RLP42 recognizes fungal endopolygalacturonases (PG) eptitope pg9(At) derived from *Botrytis cinerea*^25,26^. Similarly, RLP23 is an Arabidopsis-specific LRR-RLP, which recognizes nlp20 peptide from the NECROSIS AND ETHYLENE-INDUCING PEPTIDE1 (NEP1)-LIKE PROTEINS (NLPs) found in bacterial/fungal/oomycete species^14,27^. ReMAX is restricted to the Brassicaceae and triggered by the MAMP eMax originating from *Xanthomonas*^28^. Cuscuta Receptor1 (*CuRe1*) is specific to *Solanum lycopersicum* and senses the peptide Crip21 which originates from parasitic plants of the genus *Cuscuta*^29,30^. Finally, the inceptin receptor (INR) appears to be specific to the legume tribe Phaseoleae and recognizes inceptin (In11), a HAMP found in the oral secretion of multiple caterpillars^31–34^. Notably, all the above examples of LRR-RLPs are family-specific, restricted to the Solanaceae (Cf-2, Cf-4, Cf-9 and CuRe1), Brassicaceae (RLP23, RLP42, ReMAX) or Leguminosae (tribe Phaseoleae) (INR)^17,26^. However, despite clear signatures of lineage-specific functions, specific evolutionary steps leading to novel PRR functions across multiple species have not been described.

Mechanistic understanding of LRR-RLP sensing functions is also currently limited. No structural data exists for any PAMP or HAMP sensing LRR-RLP^26,35^. Genetic experiments using truncated or chimeric receptors have revealed subdomains essential for function of RLP23 and RLP42^26,35^, but whether similar regions mediate function for other LRR-RLPs is not clear. An alternative approach, Ancestral Sequence Reconstruction (ASR), leverages the specificity of receptor homologues for a certain PAMP/HAMP to study the emergence of a recognition function^36,37^. ASR requires dense gene trees with clearly defined sets of functional and non-functional receptors. Such data are currently unavailable for any LRR-RLP, partly due the lack of relevant high-quality genomes in closely related non-model species. Moreover, the highly duplicate-rich nature of LRR-RLP-encoding loci complicates gene annotation. However, long-read sequencing has increased the quality of *de novo* assemblies, facilitating the annotation of receptor genes at complex receptor loci^17,38,39^. Importantly, LRR-RLP recognition functions can generally be rapidly assessed through expression in a model plant, wild tobacco (*Nicotiana benthamiana*)^26,31,35,40^. A heterologous model allows rapid functional validation of potential LRR-RLP homologues, chimeric receptors and statistically inferred ancestral sequences.

In this study, we use the legume-specific LRR-RLP INR as a model to perform dense species phenotyping, comparative genomics, and functional validation, to associate gain and loss of In11 response with evolution of the contiguous *INR* receptor locus. By leveraging both existing high-quality assemblies and long-read sequencing in key legume species, we were able to study the evolution of the *INR* locus. We show that In11 response is restricted to species in the clade of the Phaseoloid legumes, which includes the agriculturally important subtribes Phaseolinae, Glycininae and Cajaninae. The contiguous receptor locus which includes *INR* is dynamic and predates the evolution of Phaseoloids, but the presence of an *INR* homologue strictly corresponds with the recognition of In11. Finally, we used chimeric and ancestrally reconstructed LRR-RLPs to gain insight into the key domains and amino acids (AA) involved in In11 recognition. We present a conceptual model for the evolution of LRR-RLP function based on patterns of INR variation in legumes.

## Results

### In11 perception is restricted to certain legume species within the Phaseoloid clade

In11 response was previously thought to be restricted to a subtribe of the legume family, the Phaseolinae, which includes cowpea (*Vigna unguiculata*) and common bean (*Phaseolus vulgaris*) but excludes soybean (*Glycine max*)^31^. To understand emergence of INR function within legumes with a higher precision, we measured ethylene accumulation triggered by In11 as a defense marker across a set of twenty-two legume species of the NPAAA papilionoids (Fig. 1a Suppl. Table 1, Suppl. Table 2). Response to In11 was only observed in plant species within the monophyletic subclade of the Phaseoloids. Hence, phylogenetic evidence suggests a single origin of In11 response at the base of the Phaseoloid legumes ca 28 mya (Fig. 1b, ★). Several of the tested plant species/accessions within this clade were unable to respond to In11, namely *Hylodesmum podocarpum*, winged bean (*Psophocarpus tetragonolobus*), *G. max*, yam bean (*Pachyrhizus erosus*) and calopo (*Calopogonium mucunoides*), although they were able to respond to the unrelated bacterial elicitor flg22 (Suppl. Fig. 1). These observations suggest the occurrence of multiple independent losses of In11 response throughout Phaseoloid evolution after the initial emergence of this function.

**Fig. 1:**
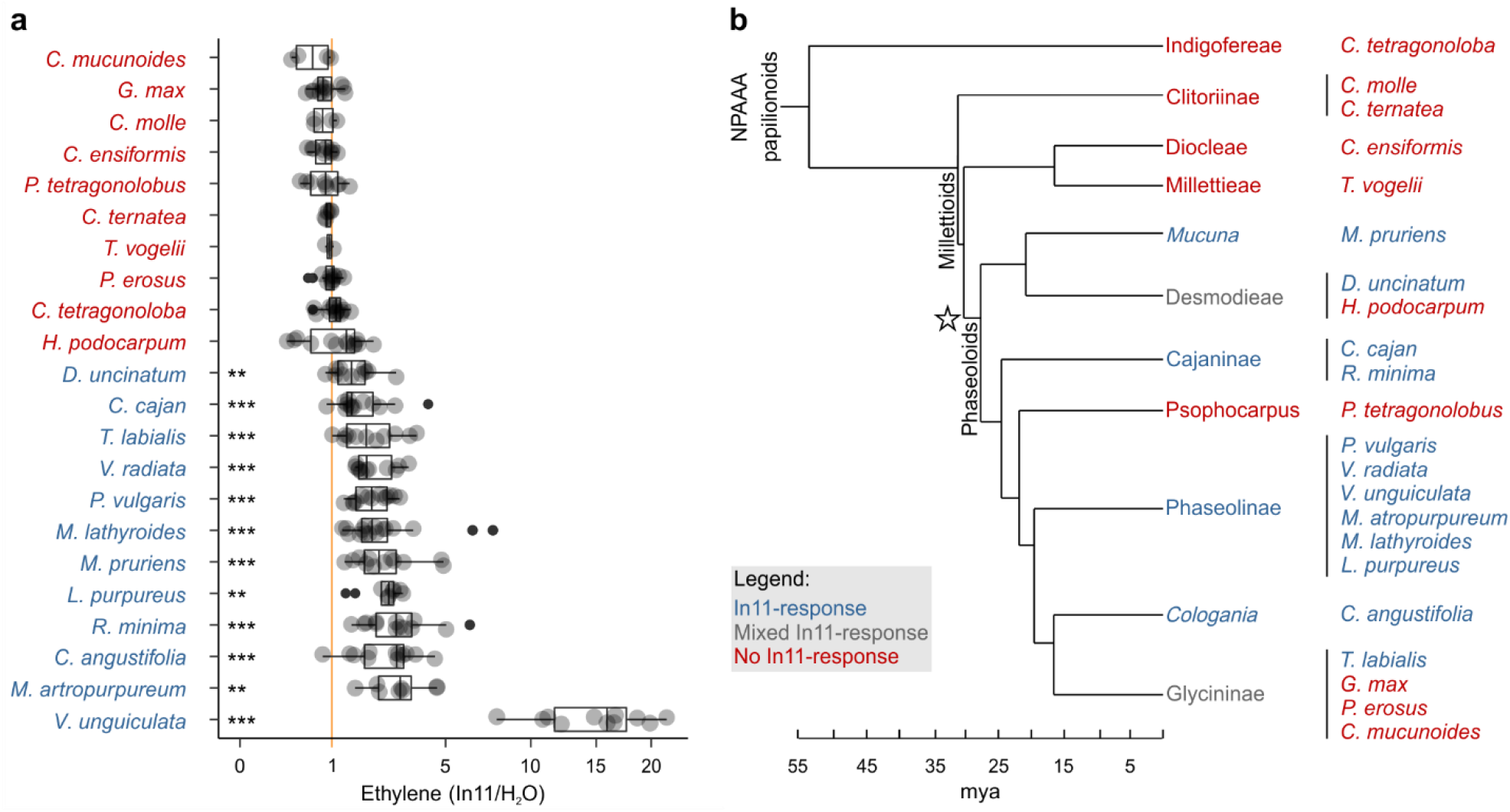
Induced ethylene response to In11 is limited to Phaseoloid legumes. **A)** Individual trifoliate leaflets were scratch wounded and treated with 1 μM In11 or H_2_O. The ratio of ethylene production for leaflets within the same leaf is shown (x-axis). The vertical orange line shows a ratio equal to one, *i*.*e*. no In11-induced ethylene burst. Biologically replicated plants are shown as separate dots. Significant differences between the control and the treatment of interest are indicated (paired Wilcoxon signed-rank test; ns non-significant, * p ≤ 0.05, ** p ≤ 0.01 and *** p ≤ 0.001). Plant species names are colored blue (significant response to In11) or red (insignificant response). Species/accessions and resulting response data can respectively be found in Suppl. Table 1 and Suppl. Table 2. **B)** A summary chronogram representing time-based phylogenetic relationships within the tested lineages, with colors representing In11 response phenotypes as in (A). The star symbol indicates the node containing all In11-responsive species at the base of Phaseoloid legumes. Divergence time is shown in million years ago (mya) and represents a composite average^83–86^.

### The contiguous *INR* locus shows LRR-RLP copy number variation in the Millettioids

To explore gain and loss of In11 response in relation to its defined PRR, we analyzed the evolution of its cognate receptor *INR*, and its genomic locus, in existing reference genomes and high quality *de novo* genome assemblies of key Phaseoloid and non-Phaseoloid species. We focused *de novo* sequencing efforts on nodes separating In11-responsive and unresponsive species in the phylogeny for comparative genomic analysis (Fig. 1b). The contiguous *INR* locus was extracted from sixteen existing legume genomes and four *de novo* assemblies: *P. erosus*, guar bean (*Cyamopsis tetragonoloba*), jack bean (*Canavalia ensiformis*) and *H. podocarpum*) (Fig. 2, Suppl. Table 3)^31^. For *de novo* assemblies obtained in this study, a combination of Nanopore and Illumina sequence data enabled the assembly of a contiguous *INR* locus in all sequenced species (Suppl. Table 4).

**Fig. 2:**
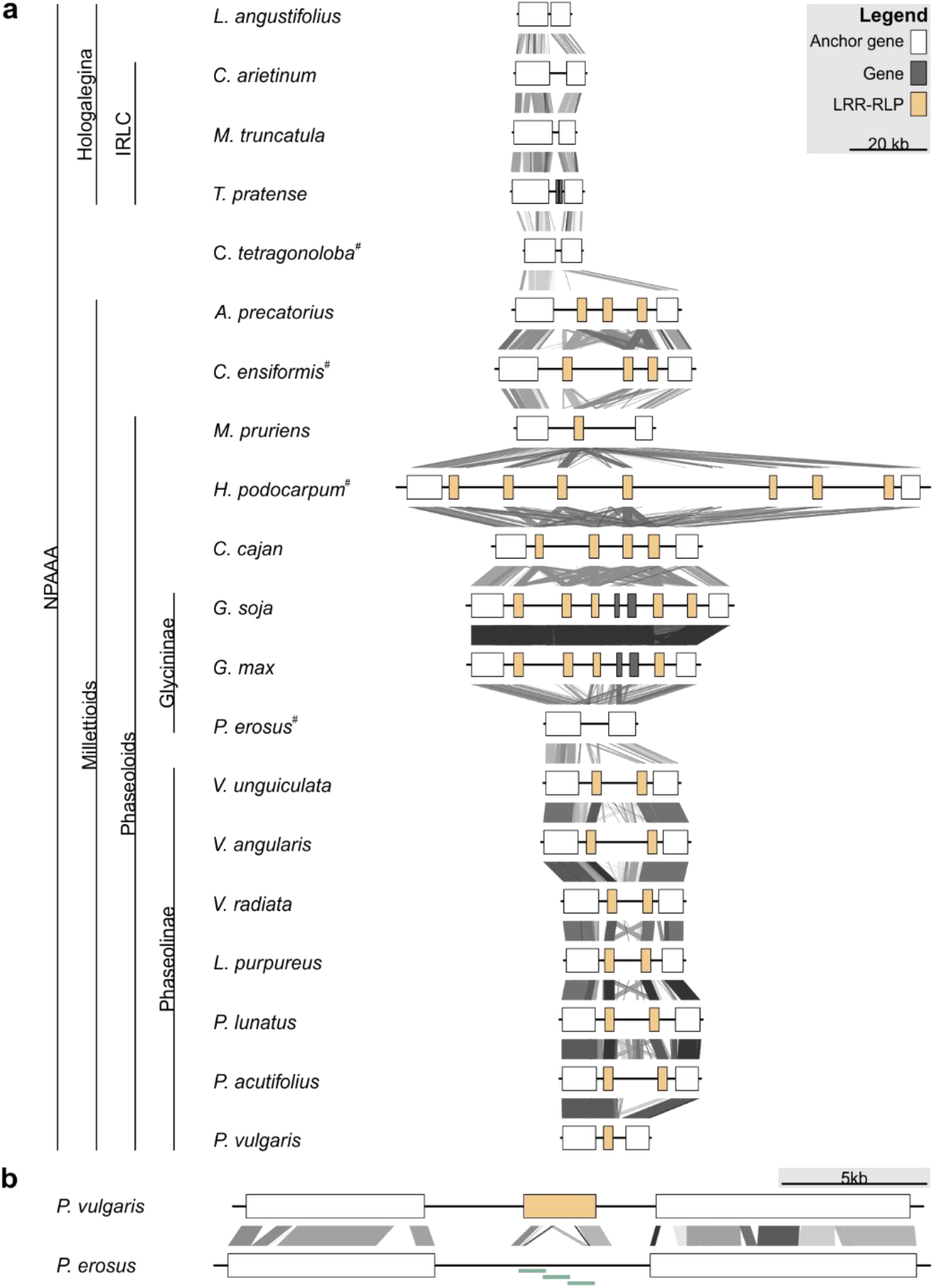
LRR-RLP copy number variation at the *INR* locus in Millettioid and non-Millettioid legume genomes. Anchor, LRR-RLP, and other genes are colored as per legend. **A)** Locus comparison of the contiguous *INR* locus of twenty NPAAA papilionoid species. Blast hits between loci are indicated with lines (e-value < 1e-04) with score according to grayscale gradient, with darker grays indicating higher similarity. Genes are labeled LRR-RLP if a complete coding sequence is present (≥ 875 AA). Species names followed by superscript (#) were newly sequenced and assembled for this project. **B)** Locus comparison of the contiguous *INR* locus of *P. vulgaris* and *P. erosus. P. vulgaris* has a functionally validated INR homologue (LRR-RLP)^31^, while *P. erosus* lacks a full-length LRR-RLP. The disruption of the *P. erosus* LRR-RLP was validated by PCR followed by Sanger sequencing as indicated with light green bars.

The organization of the *INR* locus is highly diverse among legumes (Fig. 2a), with zero to seven LRR-RLP encoding-genes per species. For all Millettioid species, BLASTN search did not identify any regions with higher score than sequences at the contiguous *INR* locus of the respective species, strongly suggesting that all potential *INR* homologues are encoded at this locus. Legume species within the Hologalegina, *Lotus angustifolius*, chickpea (*Cicer arietinum*), *Medicago truncatula* and clover (*Trifolium pratense*), do not contain an LRR-RLP encoding gene at the locus, whereas all species within the Millettioids except for *P. erosus* contained at least one LRR-RLP, consistent with a gene insertion event of an LRR-RLP at the *INR* locus in the ancestor of extant Millettioids (Fig. 1a). To investigate this emergence, we analyzed the *de novo* assembly of *C. tetragonoloba*, a close outgroup of Millettioid legumes (Fig. 1b)^41,42^. As with all species within the outgroup Hologalegina, no LRR-RLP was present between the conserved neighboring genes (i.e. anchor genes), nor did we find an LRR-RLP with greater than 68% similarity in the whole genome, strengthening support for a single ancestral RLP gene insertion event ca 32 mya (Fig. 2a).

In contrast to other closely related legume species, BLASTN analysis of the contiguous *INR* locus of *P. erosus* revealed only partial coding sequence fragments with homology to LRR-RLPs, and consequently the absence of an *INR* homologue (Fig. 2b). The *de novo* assembly of the locus was validated by performing PCR spanning the complete disruption and Sanger sequencing of the resulting amplicon. The absence of an *INR* homologue in *P. erosus* corresponds with species phenotype, namely the lack of induced ethylene response after In11 treatment (Fig. 1a).

### In11-induced functions are conferred by a single clade of LRR-RLPs (*INR* clade)

To associate *INR* locus variation with the variable In11 responses (Fig. 1), we next investigated the function and relationship of individual LRR-RLP homologues at the contiguous *INR* locus. We performed a maximum-likelihood phylogenetic analysis on the protein sequences of the LRR-RLPs within the locus across sixteen Millettioid species (Fig. 2). This analysis was supplemented with the closest related LRR-RLP genes outside the contiguous *INR* locus from *V. unguiculata* and *P. vulgaris*: *Phvul*.*007g246600* and *Vigun07g039700* (Suppl. File 2). A well-supported clade which includes the previously characterized functional *INR* from *V. unguiculata* (*Vigun07g219600*) also contained a single ortholog exclusively from plant species able to respond to In11 (Fig. 3)^31^. Hence, we hypothesized that this clade contains functional *INR* homologues which can confer In11-induced functions. To validate the putative *INR* clade, five genes (*Vigun07g219600, Phvul*.*007G077500, Mlathy INR, C*.*cajan_07316* and *Mprur INR*) within this clade were cloned and transiently expressed in the non-legume model species *N. benthamiana*. In11-induced ethylene and reactive oxygen species (ROS) production were able to be conferred by each gene (Suppl. Fig. 3), consistent with a conserved INR function in the *INR* clade (Fig. 3, blue labels).

**Fig. 3:**
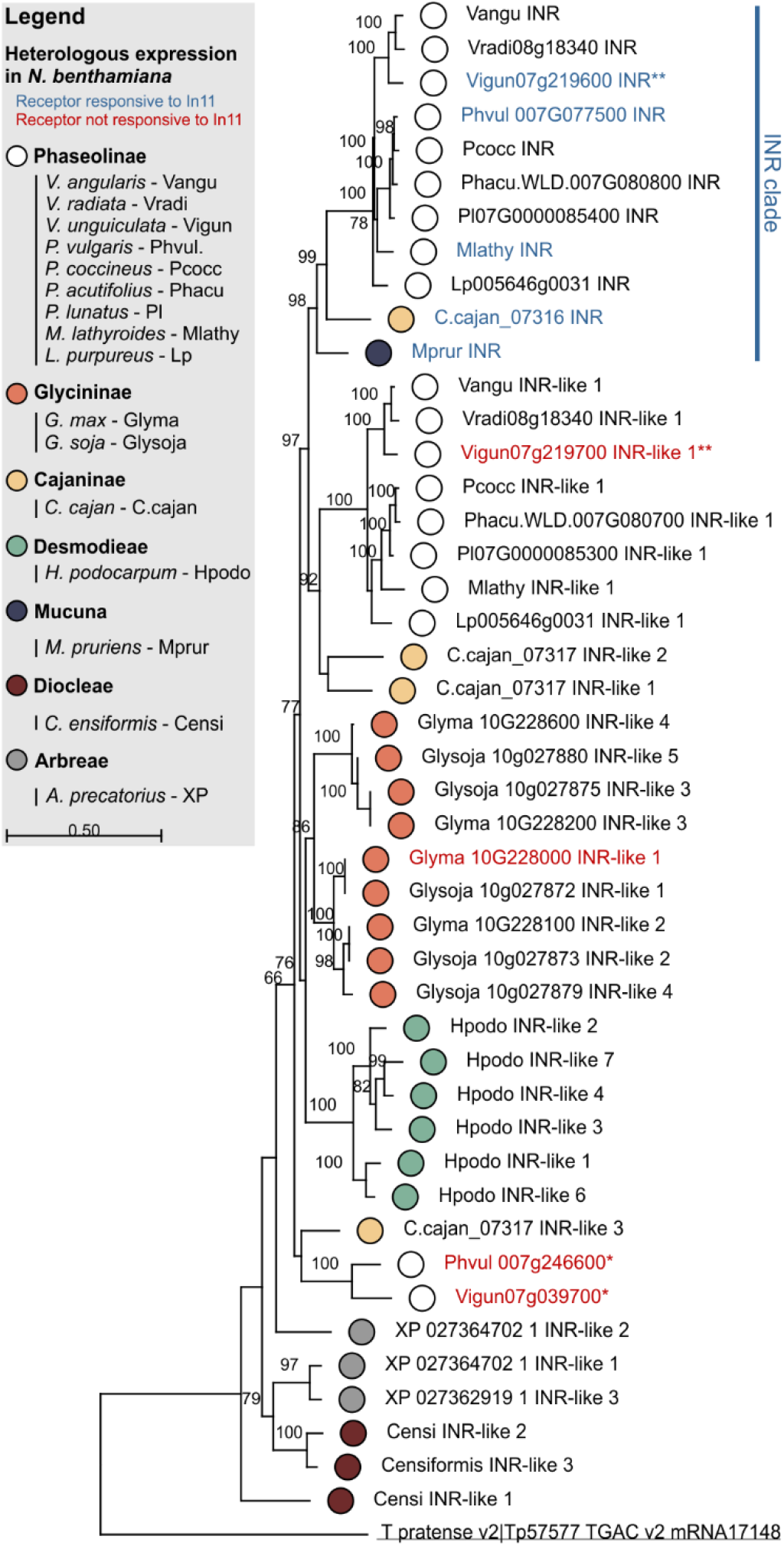
Phylogenetic analysis of LRR-RLPs at the contiguous *INR* locus reveals a clade of functional receptors. LRR-RLPs from sixteen Millettioid species are shown. Maximum likelihood analysis bootstrap values are indicated, only values higher than 65 are shown. The scale bar represents 0.5 AA substitutions per site. Filled dots indicate species of origin according to legend, where different colors indicate different subtribes. A *T. pratense* LRR-RLP was used as an outgroup to root the phylogenetic gene tree and is underlined. The functionally validated INRs of *P. vulgaris, V. unguiculata, M. lathyroides, C. cajan* and *M. pruriens* are highlighted in blue as they confer induced ROS and ethylene functions in response to In11 upon heterologous expression in *N. benthamiana* (Suppl. Fig. 3). These five validated INRs fall within the labelled “INR clade”. Heterologously expressed receptors which were not responsive to In11 are highlighted in red (Suppl. Fig. 3). One asterisk (*) highlights the LRR-RLPs which are not part of a contiguous *INR* locus. Two asterisks (**) highlight the LRR-RLPs used to create the chimeric receptors.

To assess if more distantly related, INR-like homologues could also confer In11 recognition, we measured ethylene and ROS production upon expression of LRR-RLPs outside the *INR* clade. Intriguingly, the sister clade to the *INR* clade contains LRR-RLPs from 9 out of 11 species which also have a predicted *INR* homologue in the *INR* clade itself (Fig. 3). Within this clade, we cloned and transiently expressed the LRR-RLP of *V. unguiculata* (*Vigun07g219700*). Except for *P. vulgaris* G19833, all studied Phaseolinae have an LRR-RLP in the two clades discussed above (Fig. 2). The remaining species, *G. max, Glycine soja, H. podocarpum, C. ensiformis* and *Abrus precatorius*, were In11-unresponsive (Fig. 1) and solely encode LRR-RLP receptors which fall outside the *INR* clade and its sister clade. From this group we cloned and transiently expressed the soybean LRR-RLP (*Glyma*.*10G228000*). Finally, we tested the most closely related LRR-RLP to *INR* outside the contiguous *INR* locus for both *V. unguiculata* and *P. vulgaris* (*Vigun07g039700* and *Phvul*.*007g246600*). No In11-induced responses could be observed upon heterologous expression of *Vigun07g219700, Glyma*.*10G228000, Vigun07g039700* and *Phvul*.*007g246600* (Fig. 3 and Suppl. Fig. 3). Proteins of these non-responsive LRR-RLPs were similarly or more strongly expressed in *N. benthamiana* relative to the lowest expressed responsive LRR-RLP (C.cajan_07316 INR) (Suppl. Fig. 4A), except for the marginally lower expressed Vigun07g039700, which contrasts with its closest related gene in *P. vulgaris* (*Phvul*.*007g246600*, 87% AA similarity). Thus, the ability to confer In11-induced ROS and ethylene production is strictly limited to members of the *INR* clade.

We identified and dated potential gene duplications and losses at the contiguous *INR* locus throughout its evolution within the Millettioids using NOTUNG (Suppl. Fig. 2)^43^. Reconciliation analysis between gene and species trees revealed a complex evolutionary history comprising 19 gene duplications and 20 gene losses in total. Within the Millettioids, a single duplication event gave rise to the *INR* clade containing all *INR* homologues and its sister clade which contains the closest related *INR*-like homologues. *P. vulgaris* and *Mucuna pruriens* contain an *INR* clade homologue but not members of the sister *INR*-like 1 clade, consistent with two independent gene losses. In contrast, the analysis predicts the ancestral loss of an *INR* homologue within the *INR* clade for *G. soya, G. max* and *H. podocarpum* and clade specific duplication events resulting in LRR-RLP expansions of 4-7 tandem duplicates. To confirm that *INR* was lost in *Glycine* and not just in reference assemblies, we performed BLASTP searches using Vigun07g219700 AA sequence against 26 *G. soja* and *G. max de novo* assemblies from a recent pangenome analysis^44^. Like the *Glycine* reference genomes, no BLASTP hits to Vu07g219600 for any of the 26 de novo assemblies were identified that had an AA similarity higher than 76%, suggesting that INR was lost before the speciation of *G. max* and *G. soja, i*.*e*. prior to soybean domestication.

### The C1 and C2 subdomain of the LRR ectodomain mediate In11-induced functions

To understand receptor subdomains contributing to the functional INR clade, we assembled nine chimeric receptors combining *Vigun07g219600* (*Vu*INR hereafter) and paralogous *Vigun07g219700* (*Vu*INR-like, 72% AA similarity) (Suppl. File 3). Both genes contain a typical LRR-RLP extracellular domain with 29 LRRs interrupted by a 14-AA intervening motif (C2 domain) (Fig. 4)^31^. Chimeric receptors were expressed at a similar level in *N. benthamiana* and their response to In11 was assessed by quantifying peptide-induced ethylene and ROS (Fig. 4, Suppl. Fig. 4b). Chimeric receptors with the *Vu*INR-like C1 domain were not responsive to In11. In contrast, all chimeric receptors containing the C1-C2 of *Vu*INR responded to In11 treatment with both an ethylene and ROS burst (219600-F, 219600-C3, 219600-C2 and C1-219600, Fig. 4a). Intriguingly, the chimeric receptor 219600−C1, containing C1 of *Vu*INR and C2 of *Vu*INR-like, resulted in In11-independent autoactivity. Additionally, the chimeric receptor 219600−C2, responds to In11 but has a delayed ROS burst relative to all other In11-responsive constructs tested. In summary, the *Vu*INR LRR subdomains C1 and C2 are both required for In11 response in chimeric receptors.

**Fig. 4:**
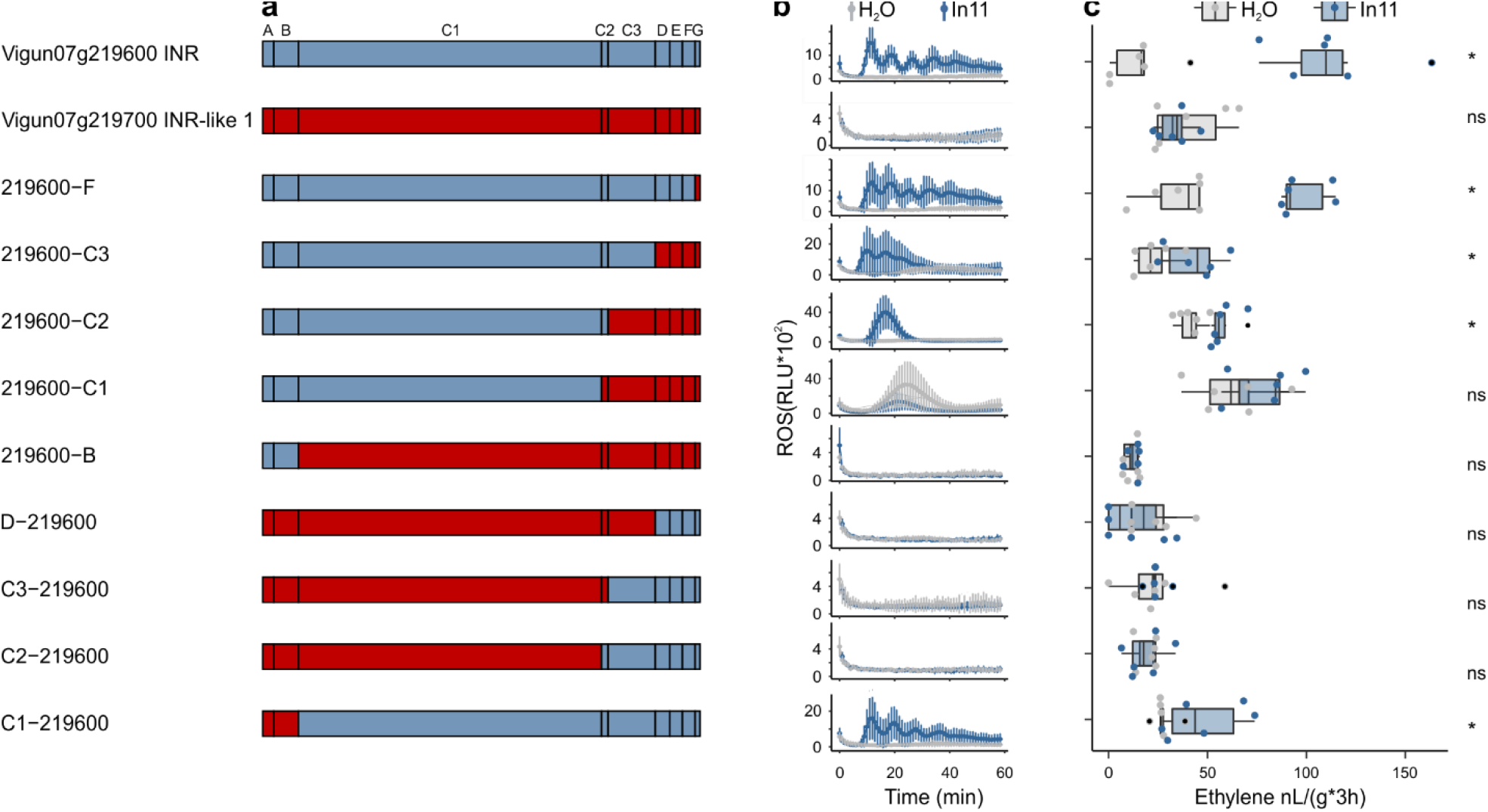
Chimeric receptors indicate that the C1 and C2 subdomains mediate INR recognition function. **A)** Schematic representation of *Vigun07g219600* (blue, *Vu*INR), *Vigun07g219700* (red, *Vu*INR-like) and the nine created chimeric receptors used for structure-function analysis. Subdomains were earlier described by Fritz-Laylin *et al*. 2005; A: putative signal peptide, B: one or two pairs of Cys that may play structural roles, C: multiple LRRs with an intervening motif (C2) inserted, D: linker domain, E: acidic domain, F: transmembrane helix, and G: cytoplasmic tail. Nucleotide sequences of the chimeric receptors can be found in Suppl. File 3. **B)** In11-dependent ROS production following the heterologous expression of receptors in *N. benthamiana*. Shown are relative luminescence units (RLU) after treatment with H_2_O (grey), or the peptide In11 (1 μM, blue Curves indicate mean +/- SD for four independent biological replicates (n=4 plants), with each biological replicate representing six technical replicates. **C)** Ethylene production following the heterologous expression of receptors in *N. benthamiana*. Ethylene production was quantified after infiltration with H_2_O (grey) or the peptide In11 (1 μM, blue). Dots represent independent biological replicates (n=6 plants). Significance was tested by performing a paired Wilcoxon signed-rank test (ns non-significant, * p ≤ 0.05).

### Ancestral sequence reconstruction (ASR) of the *INR* LRR domain

To further understand the molecular basis for functional divergence between INR and unresponsive INR-like homologues, we predicted multiple ancestral sequences of the monophyletic In11-responsive *INR* clade and its non-responsive *INR*-like 1 sister clade (Fig.5, Suppl. Fig.5, Suppl. File 4). To perform ASR, we first confirmed that nodes of interest for reconstruction were well supported by both neighbor-joining (NJ) and maximum likelihood (ML) gene phylogenies (Suppl. Fig. 5). Subsequently, we reconstructed the ancestral sequences for the LRR domain using FastML (Suppl. File 4)^45^, synthesized their predicted sequences and ligated the resulting LRR domains with flanking domains (A-B and D-G) from *Vu*INR to complete the ancestral receptor constructs (Fig. 5b). Protein expression in *N. benthamiana* was similar across all ASR variants (Suppl. Fig. 4c).

**Fig. 5:**
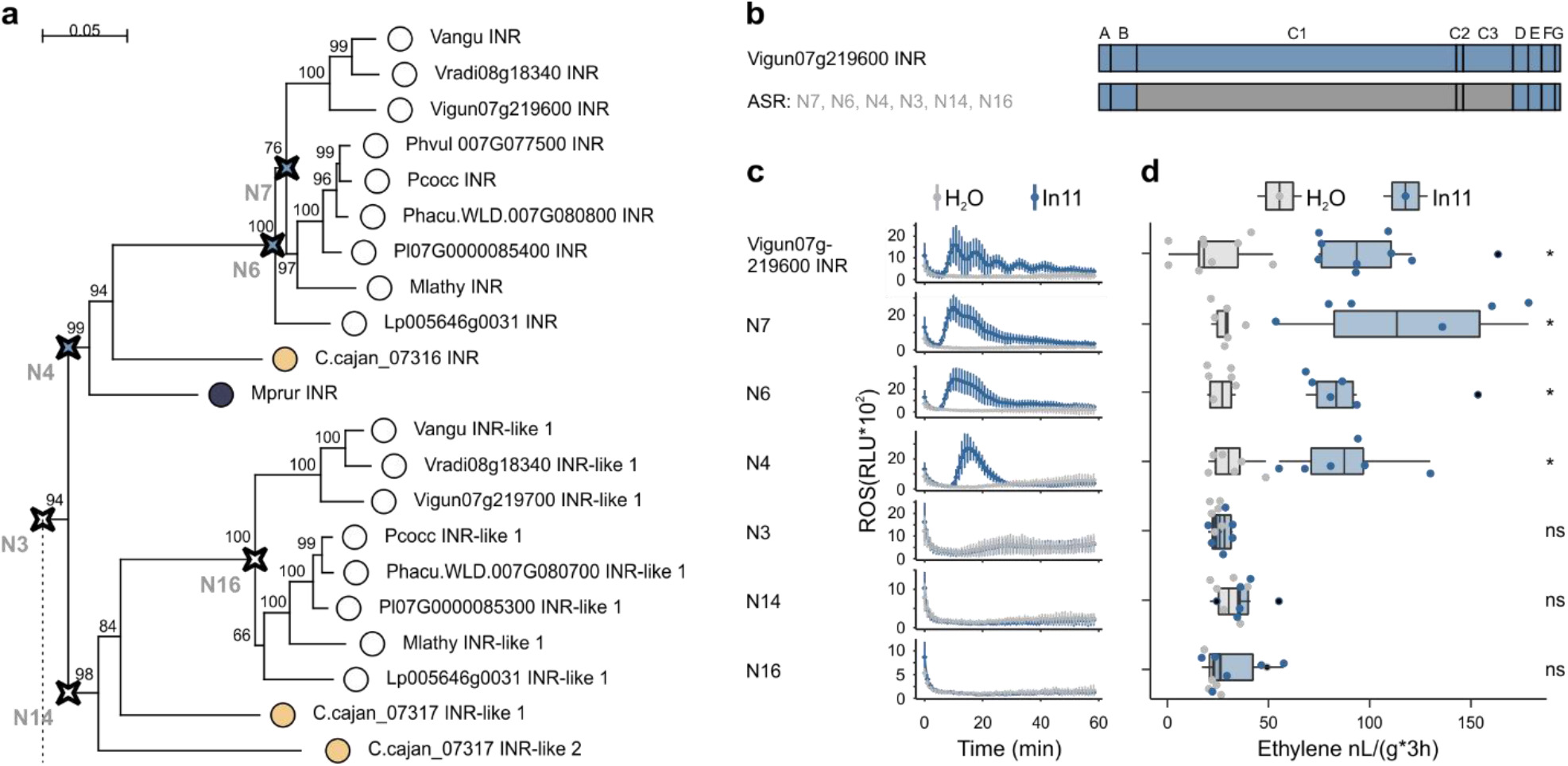
Functional analysis of ancestrally reconstructed LRR domains of INR and INR-like receptors. **A)** Part of the phylogenetic analyses of the LRR (C1-3 domain) of LRR-RLPs from the contiguous *INR* locus used for the ASR, full figure can be found in Suppl. Fig. 5. Ancestrally reconstructed nodes are marked with a compass star (N7, N6, N4, N3, N14 and N16). The scale bar represents 0.05 AA substitutions per site. Nucleotide sequences of the LRR domain can be found in Suppl. File 4. **B)** Schematic representation of *Vigun07g219600* (blue, *Vu*INR) and the six created ASR receptors used for structure-function analysis. Subdomains were earlier described by Fritz-Laylin *et al*. 2005; A: putative signal peptide, B: one or two pairs of Cys that may play structural roles, C: multiple LRRs with an intervening motif (C2) inserted, D: linker domain, E: acidic domain, F: transmembrane helix, and G: cytoplasmic tail. Nucleotide sequences of the chimeric receptors can be found in Suppl. File 3. **C)** In11-dependent ROS production following the heterologous expression of the ASR receptors in *N. benthamiana*. Relative luminescence units (RLU) are shown after treatment with H_2_O (grey), or the peptide In11 (1 μM, blue). Curves indicate mean +/- SD for four independent biological replicates (n=4 plants), with each biological replicate representing six technical replicates. **D)** Ethylene production following the heterologous expression of receptors in *N. benthamiana*. Ethylene production was quantified after infiltration with H_2_O (grey) or the peptide In11 (1 μM, blue). Dots represent independent biological replicates (n≥6 plants). Significance was tested by performing a paired Wilcoxon signed-rank test (ns non-significant, * p ≤ 0.05).

Ancestral receptors to the *INR* clade of the *Vigna, Phaseolus* and *Macroptylium* (N7), the Phaseolinae (N6) and the Phaseoloids (N4) conferred In11-induced ethylene and ROS response. In contrast, ancestral receptors of the *INR*-like 1 clade of the Phaseolinae (N16) and the Phaseoloids (N14) were not responsive to In11 (Fig. 5d, Fig. 5c). Moreover, the common ancestral receptor of the *INR* and its *INR*-like 1 sister clade (N3) is not responsive to In11 (Fig. 5). Hence, a small number of differences, 16 of 720 AA, between the LRR domain of N3 and N4, mediate differential In11 response. Moreover, only a subset of those AA is conserved within all LRR-RLPs of the *INR* clade and absent in all LRR-RLPs of the *INR*-like 1 sister clade (Suppl. Fig. 6).

## Discussion

Here we described the evolution of the legume-specific LRR-RLP INR as a functional and comparative genomic model to study A) responsiveness of an LRR-RLP against a defined elicitor and B) the evolution of novel PRR functions. We identified plant species which contain or lack a functional INR homologue, corresponding with response to the In11 HAMP elicitor. We present a conceptual model for the evolution of a novel LRR-RLP function based on patterns of *INR* locus variation in legumes. Finally, comparisons between In11-responsive and unresponsive homologues, especially through chimeric receptors and ASR, resolve potential key AA residues which mediate peptide recognition and response. Our work illuminates themes in the evolution of lineage-specific immune sensing functions, which will inform the broad use of PRRs as resistance traits^46–49^.

Evolutionary analysis of diverse plant phenotypic responses to a single PAMP or HAMP is rarely performed, although this can be a powerful approach to understand the emergence of specific immune receptor functions^2^. Patterns of In11 responses across twenty-two legume species indicated that In11 response is restricted to the Phaseoloids (Fig. 1). However, within the Phaseoloids, several tested species/accessions were not able to respond to In11, suggesting multiple independent losses of INR. These phenotypic observations allowed the investigation of the evolution of a specific LRR-RLP involved in plant immunity over a long (32my), high-resolution timescale.

We complemented broad phenotypic analysis with comparative genomics of key legume species. Analysis of the *INR* receptor locus across twenty existing and newly sequenced legume genomes revealed that INR function followed an early gene insertion and diversification at the *INR* locus. Importantly, *de novo*, long-read based assemblies presented here contained contiguous *INR* loci flanked by conserved anchor genes (Fig. 2), enabling strong conclusions with respect to receptor gene content. Comparative genomic analysis can reveal complex histories of gene insertions, duplication and losses for LRR-RKs/RLPs^50,51^. The *INR* locus shows high variability and diversification among legumes of the NPAAA papilionoids, with zero to seven LRR-RLP encoding-genes per species. Legume species within the Hologalegina (*L. angustifolius, C. arietinum, M. truncatula* and *T. pratense*) do not contain an LRR-RLP encoding gene at the locus, whereas all species investigated herein the Millettioids except for *P. erosus* contain at least one LRR-RLP (Fig. 2). This observation is consistent with a gene insertion event at the *INR* locus in the ancestor of extant Millettioids ca 32 mya.

Our analysis of INR provides an example of an LRR-RLP gene family with conserved recognition function across multiple plant genera which evolved ∼28 mya. A phylogenetic analysis revealed potential INR-homologues in ten additional species as they clustered together with *V. unguiculata* INR. Orthologous LRR-RLPs of the *INR* clade from five legume genera were tested and able to induce an In11-induced ethylene and ROS burst upon heterologous expression in *N. benthamiana* (Fig.3, Suppl. Fig. 3). This includes INR homologues from *Cajanus cajan* and *M. pruriens* of the Phaseoloid clade. Certain Phaseoloids and earlier diverged non-Phaseoloids also contain INR-like homologues at the contiguous *INR* locus; thus, INR most likely arose from an existing LRR-RLP, ∼28 mya ago. This evolutionary pattern contrasts with the evolution of the *Solanum pimpinellifolium* specific LRR-RLP Cf-2 which evolved <6 mya, potentially by intergenic recombination^49^. Our analysis thus identifies an alternative mechanism whereby an ancestral gene insertion event is the likely source of extant gene and copy number variation, preceding the evolution of a specific peptide recognition function.

The abundance of closely related legume genomes facilitates gene-species tree reconciliation, suggesting mechanisms contributing to the dynamic nature of *INR* receptor loci^43,50^. Interestingly, we observed three independent cases of *INR* loss within the Phaseoloids. Two reference genomes as well as 26 *de novo* assemblies of the closely related Glycininae, *G. soya* and *G. max*, do not encode *INR*. Nevertheless, *INR* loss does not seem to predate the divergence of the Glycininae as *Teramnus labialis* is able to respond to In11 (Fig. 1), although genome data are lacking for this species. Besides reciprocal gene loss of *INR*, gene tree-species tree reconciliation also suggests the involvement of five tandem duplications of an *INR*-like gene predating radiation of the genus *Glycine* (Suppl. Fig. 2). This is consistent with whole-genome observations of preferential gene loss in tandem clusters^50^. Similarly, within the separate Desmodieae lineage, our analysis also predicts that a *H. podocarpum INR*-like gene underwent specific tandem duplication events after the loss of *INR*, as its genome contains seven *INR*-like homologues which cluster together in a phylogenetic analysis. In contrast, *P. erosus* seems to have lost both *INR* and *INR*-like as it does not contain any complete coding sequence of an LRR-RLP at the *INR* locus. Besides *P. erosus, P. vulgaris* and *M. pruriens* also lost *INR*-like. Loss of *INR* (and *INR*-like) may reflect the propensity of tandemly duplicated loci to lose functions through gene conversion, and/or reciprocal gene loss^50^.

Our high-resolution analysis of the emergence of INR provides a roadmap for understanding other lineage-specific PRRs to detect PAMPs. While most characterized LRR-RLPs are family-specific^17,26^, extensive phenotyping of an entire plant family has not yet been conducted for well-studied PAMPs such as elf18, nlp20, pg9, crip21, csp22, and xup25. Understanding the emergence of known LRR-RK/RLPs for these PAMPs, namely EFR, RLP23, RLP42, CORE, CuRE1, and XPS1 will illuminate broader mechanisms underlying the evolution of key immune receptor modules in plants^26,27,29,52,53^. LRR-RLPs often occur in complex loci with variation within and between species ^17,26^. However, a recent pangenomic analysis of Arabidopsis indicates that several PAMP sensing LRR-RLPs occur in relatively simple and conserved loci across *A. thaliana* varieties^16^. It will be interesting to see if Arabidopsis PRR functions evolved along a similar trajectory to INR, for example via ancestral duplications preceding fixation in the Arabidopsis lineage. In summary, additional case studies for specific receptors are needed to reveal broader patterns in receptor evolution.

Our functional analysis of INR and INR-like genes in a heterologous model (*N. benthamiana*) also provides a system to study LRR-RLP function. Compared to well-studied LRR-RKs such as FLAGELLIN-SENSING 2 (FLS2)^54–56^, mechanisms underlying LRR-RLP function are not well understood. Despite multiple intense attempts, and in contrast to LRR-RKs^10,56^, structural information for ligand-binding LRR-RLPs is not available, including INR^26,35^. Moreover, Wang et al. 2019 contrasted signaling and defense responses activated by FLS2 and RLP23, but broader similarities and differences between RK and RLP families remain unclear^35,57^. Consequently, we have limited mechanistic insight into LRR-RLP function as elicitor-specific PRRs^26,35^.

Chimeric receptors formed by combining *INR* and an *INR*-like paralogue revealed RLP subdomains required for In11 response. Across all chimeric receptors, those encoding the LRR (C1) and intervening motif (C2) domains of *Vu*INR responded to In11 treatment with both an ethylene and ROS burst, suggestive of crucial elements for elicitor interaction in the C1-C2 subdomain. Notably, a chimeric receptor with mixed C1 and C2 (intervening motif), 219600-C1, resulted in In11-independent autoactivity, consistent with a critical role for C1-C2 compatibility. In addition, a chimeric receptor with mixed C2 and C3 (219600-C2) had delayed In11-induced ROS burst relative to all other In11-responsive constructs tested here (Fig. 4). These findings are consistent with previous truncation and chimeric protein analyses for Arabidopsis RLP23 and RLP42. Truncations of the RLP23 ectodomain abolished function, suggesting the necessity of the entire ectodomain for elicitor binding or proper assembly of the receptor^35^. Additionally, chimeric receptors implicated the importance of RLP42 its twelve N-terminal LRRs (C1) and LRR21-LRR24 which includes the island subdomain (C2) for recognition of fungal endopolygalacturonases^26^.

Additionally, our chimeric receptor data are consistent with a vestigial role for the cytoplasmic tail of PRRs in the LRR-RLP family. Strikingly, all identified INR homologues encode a cytoplasmic tail of only 10AA, shorter relative to all identified INR-like homologues here. Nevertheless, swapping the *Vu*INR (Vigun07g219600) cytoplasmic tail to the extended version of *Vu*INR-like 1 (Vigun07g219700), 219600-F, did not affect In11-response (Fig. 4). Previously, the complete deletion of the intracellular 17-amino-acid tail of RLP23 reduced but did not abolish receptor function^35^. Additionally, a previously described chimeric swap replacing the RLP42 terminal LRR, transmembrane helix and cytoplasmic tail with the respective subdomains of a non-responsive paralogue was still responsive to the pg9(At) elicitor^26^. Intriguingly, quantitative differences were earlier reported as the C-terminally truncated RLP23 had a reduced nlp20 response, consistent with auxiliary rather than essential function for the RLP23 cytoplasmic tail^35^.

As an alternative to the use of chimeric receptors, ASR of the *INR* LRR domain revealed additional detailed insights contributing to INR function. To trace the potential evolutionary history of INR, we used ASR by leveraging *INR* and *INR*-like homologues of sixteen legume species of the Millettioids (Suppl. Fig. 5). As the chimeric receptors implicated the LRR (C1-3) ectodomain in In11 recognition, we reconstructed ancestral LRR variants in an *INR* backbone. Our analysis suggests that the common ancestral LRR ectodomain of extant INRs conferred In11 response (N4), whereas the common ancestor of both INR and INR-like (N3) did not confer In11 response (Fig. 5). N3 and N4 differ in In11 response although they only vary by 16 AA, with only some of them conserved in all extant INRs. The role of key LRR-RLP residues in ligand-specific responses is consistent with the previous analysis of Arabidopsis RLP42. Zhang et al. 2021 introduced single AA substitutions to the functional RLP42 receptor, and several were sufficient to abolish pg9(At)-induced co-receptor association and defense responses.

A similar ASR approach to ours was employed to understand the 98-AA heavy metal-associated (HMA) domain of the plant intracellular NOD-like receptor Pik-1^37^. Specific AA changes introduced into an ancestrally reconstructed backbone were sufficient to confer effector binding and immune functions. For Pik-1 HMA, structural data provided additional insight into the mechanism of effector binding in ancestral and extant proteins. Our use of an ASR approach for a relatively long LRR domain (720 AA) now demonstrates the power of dense comparative genomic analyses to also identify key residues in extant PRRs without defined binding sites. In the absence of structural data for ligand-binding LRR-RLPs, an ASR approach may be useful to identify sets of co-varying residues critical for binding and signaling functions.

## Materials and Methods

### Plant materials

Plant species and accessions used in this study are listed in Suppl. Table 1, as well as their respective providers: Phil Miklas, (US Department of Agriculture, Prosser, WA), Creighton Miller (Texas A&M University, College Station, TX), Phil Roberts (UC Riverside, CA), Timothy Close (UC Riverside, CA) and the USDA Germplasm resources Information network (GRIN). Plants were grown in the greenhouse (25°C/21°C day/night, 60%RH and 12:12 light:dark cycles) or in growth chambers (26°C/26°C day/night, 70%RH and 12:12 light:dark cycles).

### Peptide-induced ethylene production in legume species

The In11 peptide (ICDINGVCVDA) is a host derived proteolytic fragment of the ATP synthase γ-subunit (cATPC), based on the *V. unguiculata* cATPC sequence^32^. The flg22 peptide (QRLSTGSRINSAKDDAAGLQIA) originates from bacterial flagellin^58^. Both peptides were synthesized (Genscript) and reconstituted in H_2_O. Leaflets were lightly scratch wounded with a fresh razor blade to remove cuticle area, and 10 μL of H_2_O with or without peptide (1 μM) was equally spread over the wounds with a pipette tip. After 1h, leaflets were excised and placed in sealed tubes for 2h before headspace sampling (1 mL). Ethylene was measured as previously described with a gas chromatographer (HP 5890 series 2, supelco #13018-U, 80/100 Hayesep Q 3FT x 1/8IN x 2.1MM nickel) with flame ionization detection and quantified using a standard curve (Scott, 99.5% ethylene, (Cat. No 25881-U)) (Suppl. Table 2)^59^. Subsequently, R (v4.0.3) and the R-packages dplyr (v1.0.7), ggpubr (v.0.4.0) and ggplot2 (v3.3.3) were used to analyze and plot the data, statistics were performed by using the paired Wilcoxon signed-rank test^60–62^. The resulting figure was edited in Corel-DRAW Home & Student x7.

### ROS production in legumes

Leaf punches were taken with a 4-mm biopsy punch and floated in 150 μL of H_2_O using individual cells of a white 96-well white bottom plate (BRAND*plates* F pureGrade S white (REF 781665)). After overnight incubation, ROS production was measured upon addition of a 100 μL assay solution which contains 10μg/ml luminol-horseradish peroxidase (HRP), 17μg/ml luminol and the treatment (2.5 μM In11, 2.5 μM flg22 or H_2_O). Luminescence was quantified with a TECAN SPARK plate reader every minute for an hour using an integration time of 500ms. Four technical replicates were quantified for each treatment and significant differences were determined by performing a 2-group Mann-Whitney U Test between both treatments. R and the R-packages dplyr (v1.0.7), ggpubr (v.0.4.0) and ggplot2 (v3.3.3) were used to analyze and plot the data. The resulting figure was edited in Corel-DRAW Home & Student x7.

### Genome sequencing and assembly of legume species

Leaf tissue of legume plants was harvested and grounded using an N2-chilled mortar and pestle. In contrast to *P. erosus*, nuclei isolation was first performed on frozen tissue for *C. ensiformis, C. tetragonoloba* and *H. podocarpum* using the Bionano Plant Tissue Homogenization Buffer (Part number 20283). Nuclear DNA was extracted for all above mentioned species with a modified CTAB protocol as described previously^63^. Resultant High-Molecular Weight (HMW) DNA concentrations were determined by Qubit and Bioanalyzer.

A modified protocol using the Oxford Nanopore Technologies (ONT) Rapid barcoding kit (SQK-RBK004) was used for sequencing. Briefly, 27ul of ∼20 ng/μL HMW DNA was combined with 3ul of Rapid Barcoding fragmentation mix and incubated for 1min at 30°C followed by 1 min at 80°C. Ampure beads were added at a 0.7x final concentration and bead clean-ups performed as described in the Rapid barcoding kit (SQK-RBK004). All remaining steps were performed as described in the Rapid barcoding kit protocol. Each sample was sequenced on a single MinION or PromethION flowcell (R9.4). High-accuracy base calling was performed in real time with MinKnow (v20.10.6).

Illumina short read sequence was generated for *C. tetragonoloba* and *H. podocarpum* from the same HMW DNA used for ONT long read sequencing. Paired end 2×150 bp sequence was generated on the Illumina NovaSeq6000 platform. In addition, we generated Illumina short reads for *C. ensiformis, P. erosus* and *M. lathyroides*, with GENEWIZ. For these samples, genomic DNA was extracted using the Nucleospin Plant II kit (Macherey-Nagel). The oxford nanopore sequencing data and Illumina hiseq reads used in this study can be found in SRA under Bioproject: PRJNA820752. The final genome assemblies are available on NCBI (*P. erosus*: SUB11200654, *C. ensiformis*: SUB11200580, *H. podocarpum*: SUB11200652 and *C. tetragonoloba*: SUB11200669).

The *P. erosus* genome was assembled using SPAdes (v3.15.4) using the –meta option with both the Illumina and Nanopore readsets. Additional genome assemblies were produced with FlyE (v2.8.1), consensus was generated with three rounds of Racon (v1.3.1), and finally polished with Pilon (v1.22) three times with Illumina reads (2×150 bp). Final assembly quality was determined with assembly-stats and BUSCO (v5.3.0).

### Analysis of the contiguous *INR* locus and LRR-RLP homologues

The analyzed genome assemblies, versions and their sources for the twenty legume species included in the contiguous *INR* locus analysis can be found in Suppl. Table 3^64–71^. The nucleotide sequences of the extracted loci and their coordinates can be found respectively in Suppl. File 1 and Suppl. Table 3. All INR and INR-like AA sequences included in the phylogenetic analysis can be found in Suppl. File 2. Newly sequenced and assembled genomes were analyzed to define the contiguous *INR* locus, *INR* and *INR*-like sequences. Similarly, if not yet annotated in publicly available genomes, syntenic *INR* and *INR-like* homologues were identified. First, BLASTN (BLAST 2.9.0+, e-value 10) was used to identify the *INR* syntenic locus by mining the genomes for homologues of the strongly conserved neighbor (anchor) genes of common bean *INR* (Phvul.007g077500); Phvul.007g077400 and Phvul.007g077600. Similarly, the genomes were mined for LRR-RLPs via BLASTN approach with a default e-value of 10, with exception of the *Glycine* pangenome where a BLASTP approach was used. The strongest LRR-RLP blast hits with *INR* were consistently identified in between the conserved neighbor genes. Finally, potential *INR* and *INR*-like ORFs were determined by visual inspection in IGV (v2.10.3) and compared with the sequences of closely related annotated LRR-RLPs in other legume species. Additionally, *INR* homologues were identified in *Macroptylium lathyroides* by using an alternative method. First, *M. lathyroides* Illumina HiSeq paired end reads were mapped against both *Phvul*.*007G077500* (*INR*) and *PvUI111*.*07G078600* (*INR*-like) by using bwa (v0.7.17-r1188)^72^. Second, mapped reads were sorted and indexed by using samtools (v 1.13)^73^. Third, IGV (v2.10.3) was used to inspect the mapped reads and identify the SNPs of Mlathy *INR* and *INR*-like in comparison to the *P. vulgaris* homologues. Fourth, the above three steps were reiterated with the newly acquired gene sequences. Finally, the acquired Mlathy *INR* and *INR*-like sequences were confirmed by PCR using the Q5 Hot Start High-Fidelity kit (NEB), enzymatic clean-up using ExoSAP-IT™ (Thermo Fisher scientific) and Sanger sequencing of the entire amplified PCR product (GENEWIZ). Primers used in this process are listed in Suppl. Table 5.

### Contiguous *INR* locus analysis and validation of *P. erosus* receptor disruption

Locus comparison was performed using R (v4.0.3) and the R-package genoPlotR (v0.8.11) using the extracted contiguous *INR* loci and their corresponding annotation (Suppl. File 1). The resulting figure was edited in Corel-DRAW Home & Student x7. A PCR was performed to validate the disruption of the receptor at the syntenic locus of *P. erosus* using the Q5 Hot Start High-Fidelity kit (NEB). Primers used in the reaction are listed in Suppl. Table 5. Subsequently, the amplified PCR product was enzymatically cleaned-up using ExoSAP-IT™ (Thermo Fisher scientific). Finally, the disruption was validated by Sanger sequencing of the entire amplicon (GENEWIZ).

### Phylogenetic analysis of *INR* and *INR-like* homologues

Sequences were aligned using the online version of MAFFT 7 using the E-INS-i strategy^74^. One potential pseudogene of *H. podocarpum* was not incorporated in the phylogenetic analysis since the length of the sequence was about 79% of cowpea *INR* due to the absence of a part of the LRR domain in comparison to all other annotated receptors at the contiguous *INR* locus. A phylogenetic analysis was performed on the CIPRES web portal using RAXML-HPC2 on XSEDE (v8.2.12) with the automatic protein model assignment algorithm using maximum likelihood criterion and 250 bootstrap replicates^75,76^. The DUMMY2 protein model was selected as the best scoring model for maximum likelihood analysis. The resulting tree was rooted, visualized using MEGA10 and edited in Corel-DRAW Home & Student x7.

### Notung analysis: prediction of duplication and gene loss events at the contiguous *INR* locus

First, a phylogenetic analysis was performed similar to the approach above with the sole difference that only *INR* and *INR-*like homologues located at the contiguous *INR* locus were included. Second, a species tree in Newick format was built which includes all species of which 1) *INR* and *INR-*like homologues were extracted, 2) *C. tetragonoloba* as it is the closest related legume species without a receptor at the contiguous *INR* locus, and 3) *M. truncatula* as the LRR-RLP used as an outgroup in the analysis was extracted from its genome. Third, the NOTUNG analysis was performed using the default options, including a duplication cost of 1.5 and a loss cost of 1 (v2.9.1.5)^43^. NOTUNG ignores incomplete lineage sorting as an evolutionary mechanism when both a rooted species and gene tree are used as input, as was the case for the present study.

### Molecular cloning of *INR* and *INR-*like homologues

Leaf tissue of legume plants was harvested and grounded using an N2-chilled mortar and pestle. Genomic DNA was extracted using the Nucleospin Plant II kit (Macherey-Nagel). RNA was extracted using the Nucleospin RNA Plant kit (Macherey-Nagel). cDNA was created using SuperScript™ IV Reverse Transcriptase (Invitrogen). All constructs were created using a hierarchical modular cloning approach facilitated by the MoClo toolkit and the MoClo Plant Parts kit^77,78^. LRR-RLPs with no introns were PCR amplified from genomic DNA (Q5 Hot Start High-Fidelity, NEB), all others (*Phvul*.*007g246600* and *Vigun07g039700*) were amplified from cDNA. All primers used for amplification are listed in Suppl. Table 5. All amplified PCR fragments were gel extracted using the PureLink™ Quick Gel Extraction Kit (Thermo Fisher scientific) and purified and concentrated using the Monarch PCR & DNA Cleanup Kit. Subsequently, the PCR fragments were ligated in an L-1 acceptor vector. A second digestion/ligation step was completed, to ligate multiple parts together and complete the CDS while inserting it in an L0 acceptor vector. Throughout the previous steps, recognition sites for BsaI and/or BpiI were removed from the CDS. All constructs were validated by Sanger sequencing upon completion. Finally, all constructs were completed by adding the following MoClo modules; 35s Caulifower Mosaic Virus + 5’UTR Tobacco mosaic virus (pICH51266), GFP (*A. victoria*) (pICSL50008) and the OCS1 terminator (pICH41432)^78^.

### *N. benthamiana* transient expression and Western blotting

Constructs were electroporated into *Agrobacterium tumefaciens* strain GV3101(pMP90,pSOUP). Overnight cultures were resuspended in 150 μm acetosyringone in 10 mM 2-(*N*-morpholino) ethanesulfonic acid (MES), pH 5.6, 10 mM MgCl_2_. After 3h of incubation at room temperature, *N. benthamiana* leaves of 5-week-old plants were infiltrated at an optical density of 0.45 at 600 nm (OD_600_).

Leaf punches were taken 48h after infiltration of the *N. benthamiana* leaves to validate the expression of the constructs in *N. benthamiana* by Western blot (Suppl. Fig. 4). Ground, frozen tissue was homogenized in a 3x lamellae buffer (50 mM Tris-Cl pH 6.8, 6% SDS, 30% glycerol, 16% β-mercaptoethanol and 0.006% Bromophenol blue) and then cleared by centrifugation (10 m, 20,000 rcf). Subsequently, proteins in the supernatant were separated by performing an SDS-PAGE on an 8% acrylamide gel. Finally, a Western blot was performed to visualize the GFP-tagged heterologously expressed proteins and actin as a loading control with respectively an α-GFP polyclonal (A-6455; Thermo) primary antibody at a 1:2,000 dilution and an anti-Actin (ab197345, abcam) primary antibody at a 1:5,000 dilution. α-rabbit (A6154; Sigma) was used as a secondary antibody for both at 1:10,000 dilution.

### ROS measurements in *N. benthamiana*

Following *Agrobacterium* infiltration for receptor expression (24 h), leaf punches were taken with a 4-mm biopsy punch and floated in 150 μL of H_2_O using individual cells of a white 96-well white bottom plate (BRAND*plates* F pureGrade S white). Subsequently, the same procedure was followed as outlined for ROS production in legumes. Four biological replicates were quantified (n=4 plants), with each biological replicate representing six technical replicates. R and the R-packages dplyr (v1.0.7), ggpubr (v.0.4.0) and ggplot2 (v3.3.3) were used to analyze and plot the data. The resulting figure was edited in Corel-DRAW Home & Student x7.

### Ethylene in *N. benthamiana*

For ethylene assays in *N. benthamiana*, a fully expanded leaf of 5wk-old plants was infiltrated with H_2_O or 1μM In11 with a blunt syringe. Subsequently, four leaf discs within the infiltrated area were immediately excised with a no. 5 cork borer and sealed in tubes^31^. Headspace ethylene was measured after 3 h of accumulation as described above. Subsequently, R and the R-packages dplyr (v1.0.7), ggpubr (v.0.4.0) and ggplot2 (v3.3.3) were used to plot the data and perform the statistics (paired Wilcoxon signed-rank test). The resulting figure was edited in Corel-DRAW Home & Student x7.

### Construction of chimeric receptors

Chimeric LRR-RLP constructs were generated using the MoClo toolkit^77^. MoClo overhangs were designed in such a way that specific fragments amplified from the L0 constructs of *Vigun07g219600* and *Vigun07g219700* could be ligated in the preferred direction and order in an L0 vector. Primers used in these reactions are listed in Suppl. Table 5. All constructs were validated by Sanger sequencing upon completion. Subsequently, CDS stored in L0 universal acceptors were combined with the same MoClo modules as the earlier described LRR-RLP constructs mentioned above to create a complete L1 construct. Transient expression of chimeric receptor constructs in *N. benthamiana* was validated with western blot.

### ASR of LRR domain

The LRR (C1-3 domain, Fig. 4) were extracted from all LRR-RLP sequences included in the earlier mentioned phylogenetic analysis (Fig. 3, Suppl. File 4). The LRR domain and subdomains of *Vu*INR were earlier identified using LRRfinder^31,79^. Phylogenetic trees were built using MEGA X software^80^, and bootstrap method based on 1000 iterations. A codon-based 2172-nucleotide-long alignment was generated using MUSCLE^81^. NJ clustering method was used for constructing the codon-based tree on Maximum Composite Likelihood substitution models. The ML tree was calculated using the GTR+G+I submodel as implemented in MEGA X software^80^. The resulting ML tree was used for the ASR of selected nodes of interest. Joint and marginal ASR were performed with FastML software using Jukes and Cantor substitution model for nucleotides, gamma distribution, and 90% probability cutoff^82^. Finally, the sequences were domesticated to facilitate MoClo cloning by removing the BsaI and BpiI cut sites (Suppl. File 4).

### Construction of receptors with an ancestral reconstructed LRR domain

Ancestral reconstructed LRR-RLP constructs were generated using the MoClo toolkit^77^. MoClo overhangs were designed in such a way that the AB domain and the DEFG domain amplified from *Vigun07g219600* could be ligated with the synthesized ancestral reconstructed LRR domain (C domain), resulting in an L0 vector. Primers used in these reactions are listed in Suppl. Table 5. All constructs were validated by Sanger sequencing upon completion. Subsequently, the complete CDS stored in L0 acceptors were combined with the same MoClo modules as the earlier described LRR-RLP constructs mentioned above to create a complete L1 construct. Transient expression of the ancestral reconstructed LRR-RLPs in *N. benthamiana* was validated with western blot.

## Supporting information

Suppl. Table 1: Overview of the used legume species, accessions, and their origin

Suppl. Table 2: Overview of ethylene response data of legume species and the details of the experimental set-up

Suppl. Table 3: Overview of mined assemblies of contiguous INR loci (+ coordinates) and LRR-RLPs

Suppl. Table 4: Genome assembly stats

Suppl. Table 5: Primers used in this study

Suppl. File 1: Fasta file of nt sequences of the INR syntenic loci incorporated in the contiguous INR locus analysis

Suppl. File 2: Fasta file of the AA sequences INR and INR-like homologues included in the phylogenetic analysis

Suppl. File 3: Fasta file containing the sequences of the chimeric receptors

Suppl. File 4: Fasta file containing the nucleotide LRR domain sequences of the INR and INR-like homologues used for the ASR analysis and the resultin

## Acknowledgments

S.S. is supported as a Belgian American Educational Foundation postdoctoral fellow and the Mary Race Bevis Postdoc Research Award. S.S. and A.D.S. are supported by start-up funding from the University of Washington. A.D.S. is a Distinguished Investigator of the Washington Research Foundation.

## Contributions

S.S. and A.D.S. conceived and designed the experiments; S.S., B.W.A and A.G.K.G. conducted experiments; S.S, B.W.A., A.N.E and A.D.S. analyzed data; S.S. prepared all figures; S.S. and A.D.S. wrote the manuscript. All authors discussed the results and commented on the manuscript.

## Supplemental Figures

**Suppl. Fig. 1:**
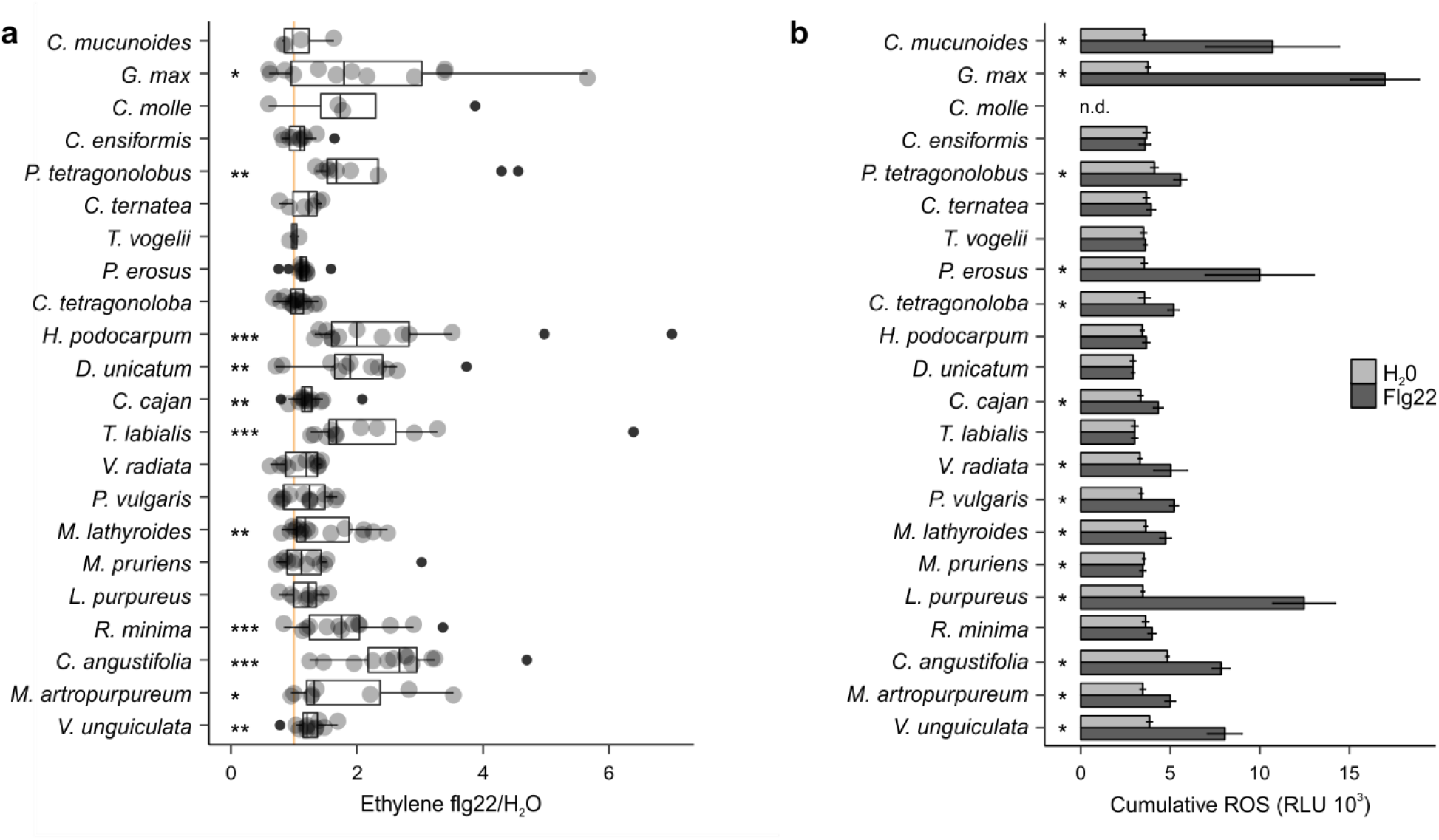
Flg22-induced ROS and ethylene responses are idiosyncratic across Millettioid and non-Millettioid legume species. **A)** Individual trifoliate leaflets were scratch wounded and treated with 1 μM flg22 or H_2_O. The ratio of ethylene production for leaflets within the same leaf is shown (x-axis). Species are listed in order of In11/H_2_O response as per Fig. 1. The vertical orange line shows a ratio equal to one, *i*.*e*. no induced ethylene response to In11. Biological replicate plants are shown as separate dots. Significant differences between the control and the treatment of interest are indicated (paired Wilcoxon signed-rank test; ns non-significant, * p ≤ 0.05, ** p ≤ 0.01 and *** p ≤ 0.001). **B)** ROS-bursts are used a second marker of plant immunity response upon application of flg22. Shown is cumulative ROS data (3-60min) of four technical replicates in relative luminescence units (RLU) over 1 hour (1 observation/minute) after treatment with H_2_O, or the peptide flg22 (1 μM). No ROS data was determined for *Centrosema molle* (n.d.). Significant differences between the control and the treatment of interest were found by performing a 2-group Mann-Whitney U Test (* p ≤ 0.05).

**Suppl. Fig. 2:**
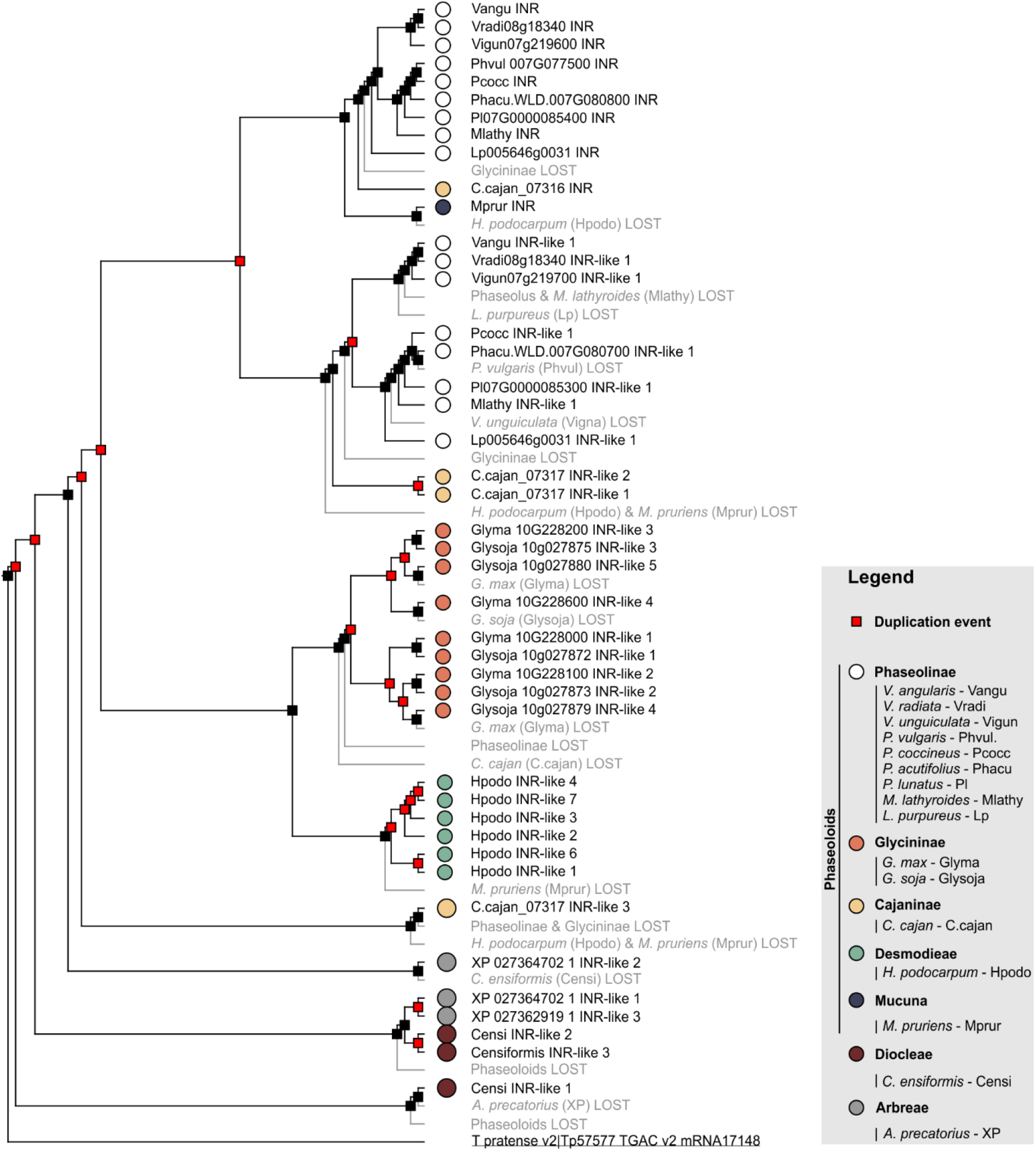
Prediction of gene duplication and gene loss events throughout the evolution of the contiguous *INR* locus in the Millettioids. Gene tree-species tree reconciliation to identify the duplication and loss events at each branch using Notung^43^. Predicted duplication events are marked with a red square, black squares represent speciation events, and lost nodes/genes are highlighted in grey. Filled dots indicate species of origin according to legend, where different colors indicate different subtribes. A *T. pratense* LRR-RLP was used as an outgroup to root the phylogenetic gene tree and is underlined.

**Suppl. Fig. 3:**
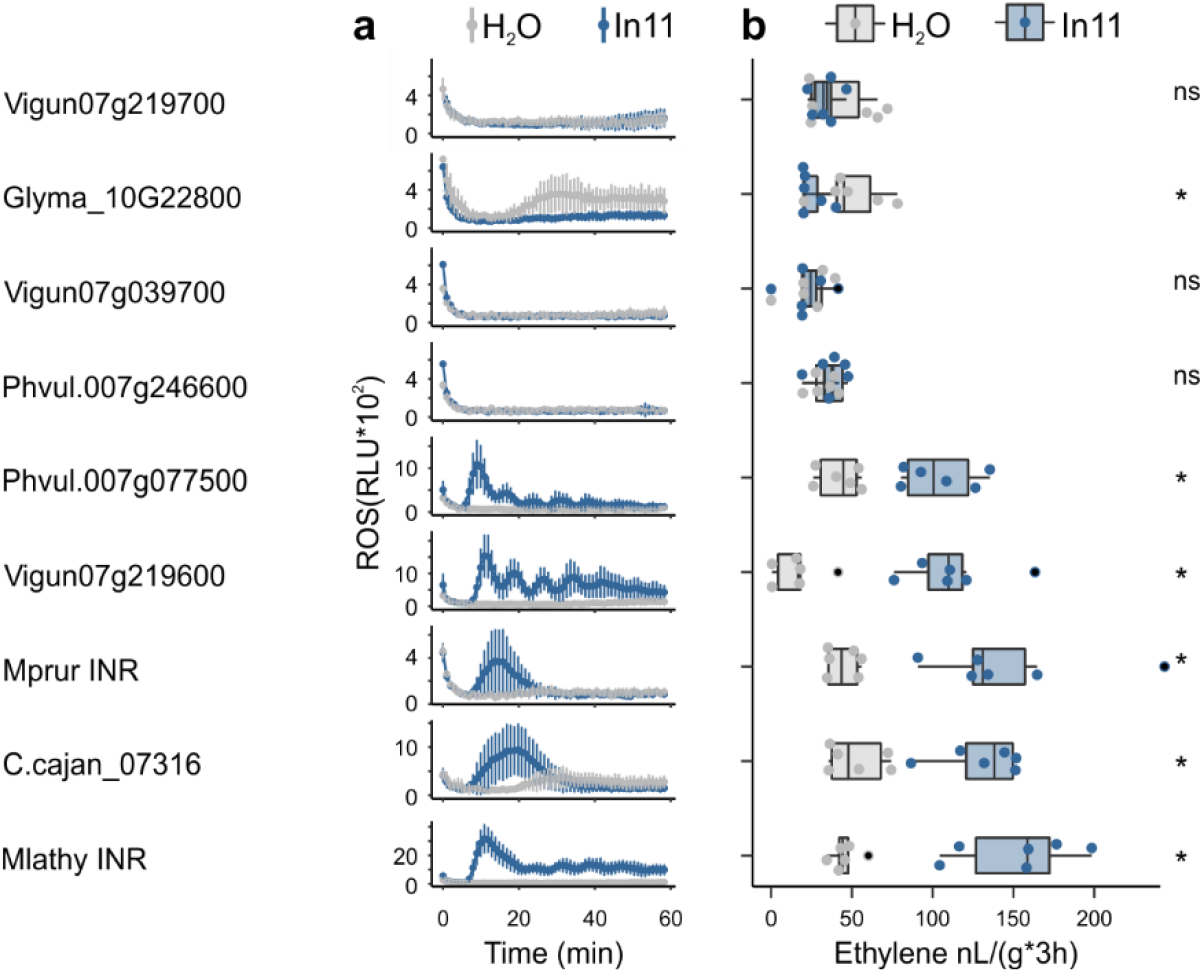
In11-induced ROS and ethylene production conferred by INR and INR-like after heterologous expression. In11-dependent plant immunity response data following heterologous expression of multiple receptor constructs in *N. benthamiana*. **A)** Shown are cumulative relative luminescence units (RLU) after treatment with H_2_O (grey), or the peptide In11 (1 μM, blue). Curves indicate mean +/- SD for four independent biological replicates (n=4 plants), with each biological replicate representing six technical replicates. **B)** The x-axis shows the amount of ethylene released after infiltration with 1 μM In11 (blue) or water (grey). Dots represent independent biological replicates (n=6 plants). Significance was tested by performing a paired Wilcoxon signed-rank test (ns non-significant, * p ≤ 0.05).

**Suppl. Fig. 4:**
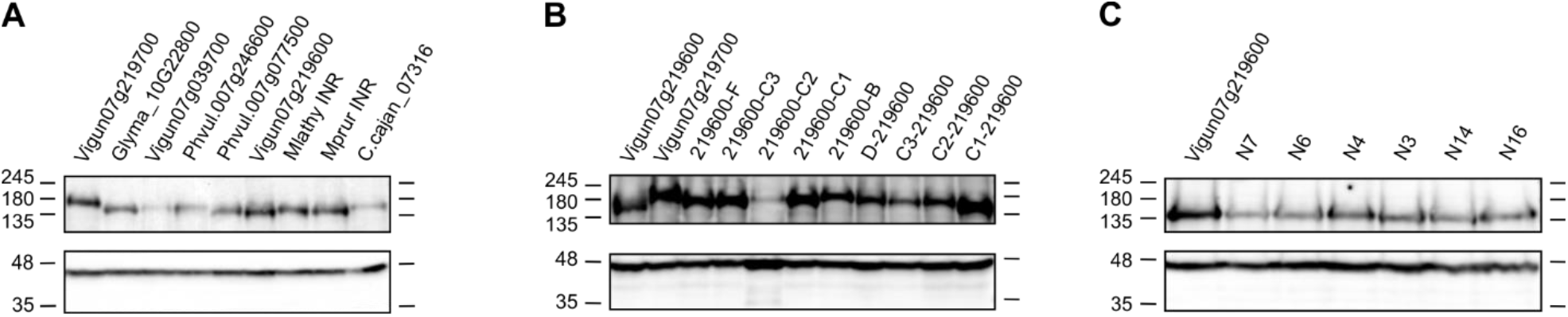
Western blot of the heterologously expressed constructs in *N. benthamiana*. Tissue was harvested 48h after construct infiltration in *N. benthamiana*. Western blots were probed with 1) GFP antibody as the receptor constructs had a C-terminal GFP tag (top), and 2) actin antibody as a loading control (bottom). Western blots show all heterelogously expressed receptors of which response to In11 was tested: **A)** INR and INR-like homologues from diverse legumes, **B)** chimeric receptors, **C)** ASR receptors.

**Suppl. Fig. 5:**
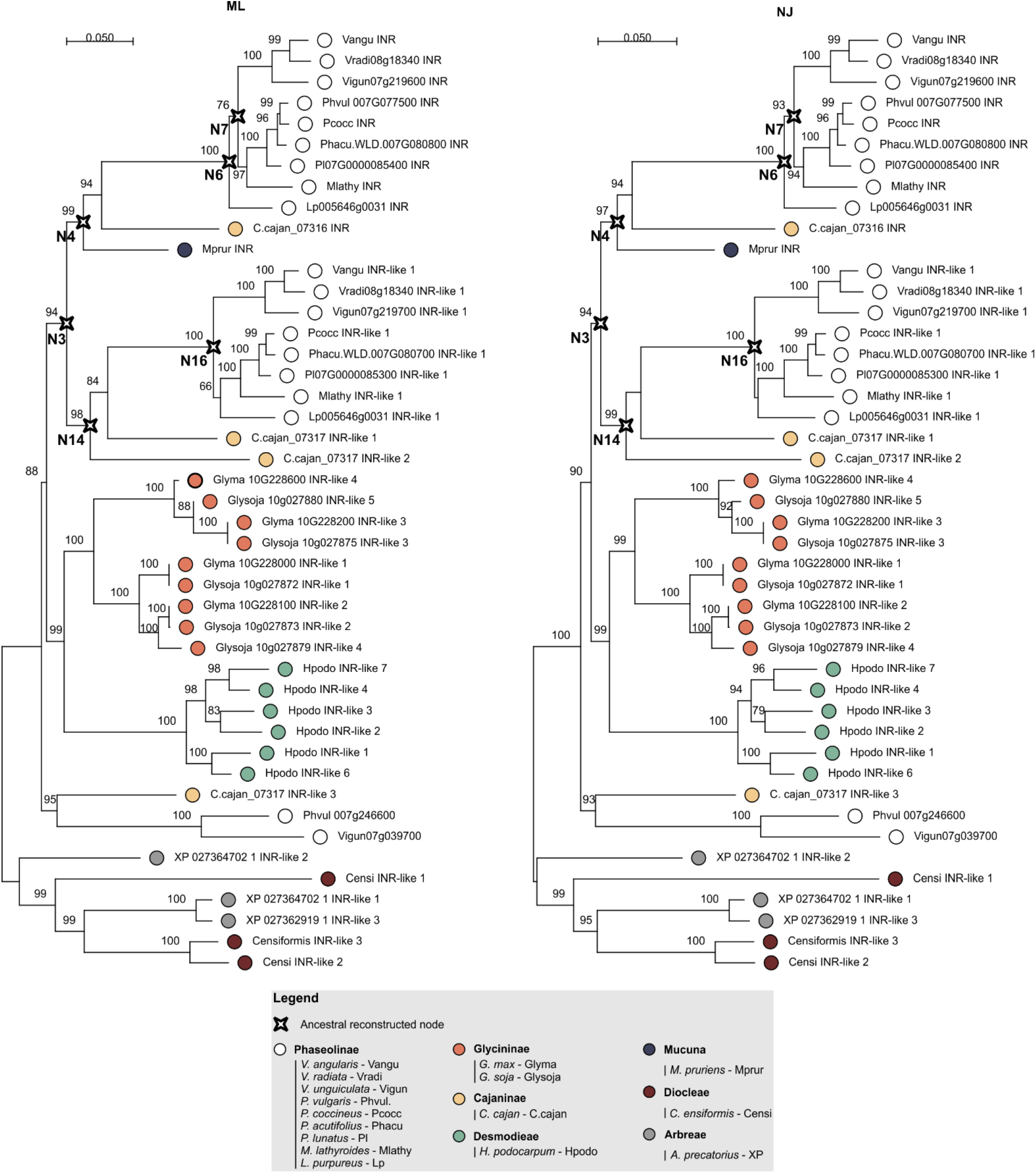
Phylogenetic analyses of the LRR (C1-3 domain) of LRR-RLPs from the contiguous *INR* locus. The phylogenetic trees were built using MEGA X software and 1000 bootstraps^80^. Maximum likelihood (ML – left side) and neighbor joining (NJ – right side) trees were calculated based on all codon positions of a codon-based alignment^81^. Maximum likelihood analysis bootstrap values are indicated, only values higher than 65 are shown. The scale bar represents 0.05 AA substitutions per site. The ancestral reconstructed nodes selected for functional validation are marked with a compass star (N7, N6, N4, N3, N14 and N16). Filled dots indicate species of origin according to legend, where different colors indicate different subtribes.

**Suppl. Fig. 6:**
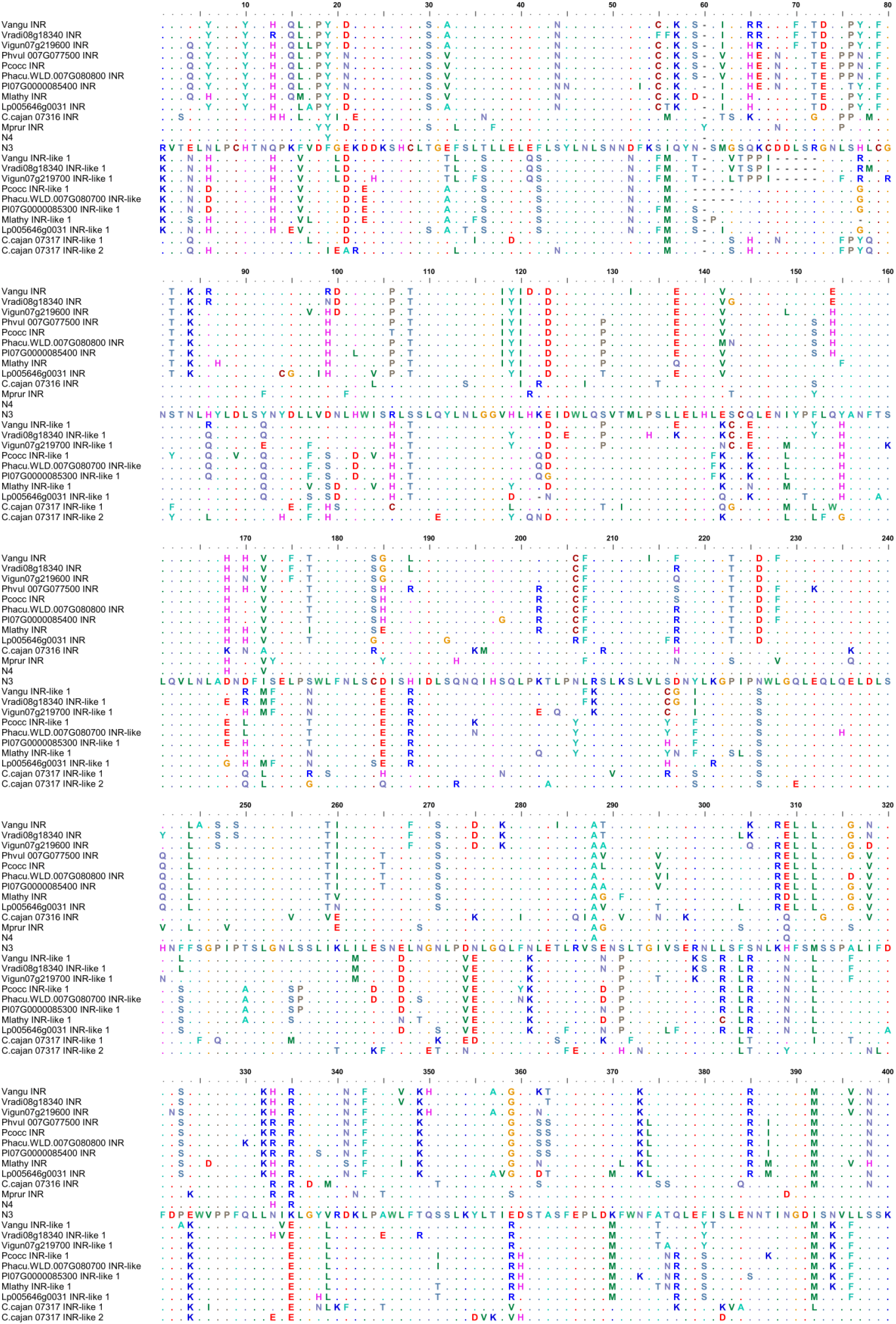
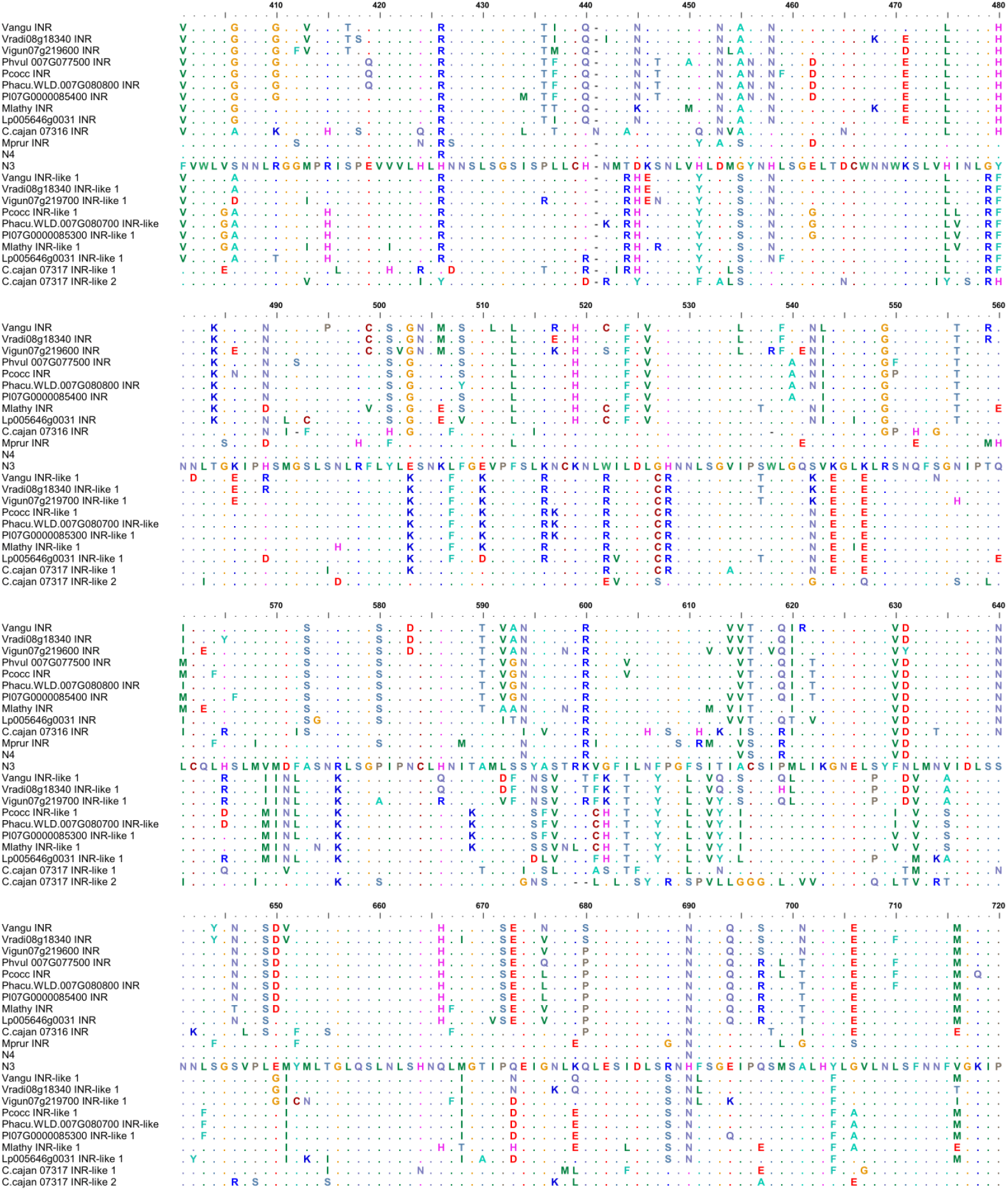
AA sequence alignment of the LRR domains of the *INR* clade, INR-like 1 clade and the closest ancestral reconstructed nodes with differential In11-response.

## Supplemental Table legends

**Suppl. Table 1: Overview of the used legume species, accessions, and their origin**

**Suppl. Table 2: Overview of ethylene response data of legume species**

**Suppl. Table 3: Overview of mined assemblies of contiguous *INR* loci (+ coordinates) and LRR-RLPs**

**Suppl. Table 4: Genome assembly stats**

**Suppl. Table 5: Primers used in this study**

## Supplemental file legends

**Suppl. File 1: Fasta file of nt sequences of the INR syntenic loci incorporated in the contiguous *INR* locus analysis**

**Suppl. File 2: Fasta file of the AA sequences INR and INR-like homologues included in the phylogenetic analysis**

**Suppl. File 3: Fasta file containing the sequences of the chimeric receptors**

**Suppl. File 4: Fasta file containing the nucleotide LRR domain sequences of the INR and INR-like homologues used for the ASR analysis and the resulting predicted (and domesticated for MoClo) LRR domain sequences for the ancestral nodes of interest**

